# Correction for participation bias in the UK Biobank reveals non-negligible impact on genetic associations and downstream analyses

**DOI:** 10.1101/2022.09.28.509845

**Authors:** Tabea Schoeler, Doug Speed, Eleonora Porcu, Nicola Pirastu, Jean-Baptiste Pingault, Zoltán Kutalik

## Abstract

While large-scale volunteer-based studies such as the UK Biobank (UKBB) have become the cornerstone of genetic epidemiology, the study participants are rarely representative of their target population.

Here, we aim to evaluate the impact of non-random participation in the UKBB, and to pin down areas of research that are particularly susceptible to biases when using non-representative samples for genome-wide discovery. By comparing 14 harmonized characteristics of the UKBB participants to that of a representative sample, we derived a model for participation probability. We then conducted inverse probability weighted genome-wide association analyses (wGWA) on 19 UKBB traits. Comparing the output obtained from wGWA (N_effective_=94,643 – 102,215) to standard GWA analyses (N=263,464 – 283,749), we assessed the impact of participation bias on three estimated quantities, namely 1) genotype-phenotype associations, 2) heritability and genetic correlation estimates and 3) exposure-outcome causal effect estimates obtained from Mendelian Randomization. Participation bias can lead to both overestimation (e.g., cancer, education) and underestimation (e.g., coffee intake, depression/anxiety) of SNP effects. Novel SNPs were identified in wGWA for 12 of the included traits, highlighting SNPs missed as a result of participation bias. While the impact of participation bias on heritability estimates was small (average change in *h*^2^: 1.5%, maximum: 5%), substantial distortions were present for genetic correlations (average absolute change in *r*_*g*_: 0.07, maximum: 0.31) and Mendelian Randomization estimates (average absolute change in standardized estimates: 0.04, maximum: 0.15), most markedly for socio-behavioural traits including education, smoking and BMI. Overall, the bias mainly affected the magnitude of effects, rather than direction. In contrast, genome-wide findings for more molecular/physical traits (e.g., LDL, SBP) exhibited less bias as a result of selective participation.

Our results highlight that participation bias can distort genomic findings obtained in non-representative samples, and we propose a viable solution to reduce such bias. Moving forward, more efforts ensuring either sample representativeness or correcting for participation bias are paramount, especially when investigating the genetic underpinnings of behaviour, lifestyles and social outcomes.

## Introduction

Elucidating the genetic underpinnings of health and disease is the overarching aim of genetic epidemiology. Fast-growing biobanks with rich phenotypic data are therefore curated, to maximize power for genome-wide discovery. To ensure validity of findings obtained from genome-wide studies, substantial efforts are made to eliminate potential sources of bias, such as population stratification, assortative mating, measurement error or indirect genetic effects^1–4^. A particularly challenging bias – and typically not considered in genetic studies – can occur when data is collected from individuals not representative of their target population^5,6^. While valid conclusions are possible in non-representative samples under certain conditions (e.g., if study participation is unrelated to both the independent and dependent variable), study participation is linked to many commonly studied factors, including mental and physical health, substance use (cigarettes, alcohol), income and educational attainment^7–11^ – where participants typically show a better health profile than the target population. Such ‘healthy-volunteer bias’ is well documented in the UK Biobank (UKBB), one of the most widely used resources for biomedical research. Of the 9 million people invited to participate in the UKBB, only 5.5% (~500,000) were recruited into the study – a sample of volunteers with more healthy lifestyles, higher levels of education, and favourable health profiles compared to the general population^12,13^.

Given the growing reliance on non-representative biobanks, it is paramount to assess the extent to which study participation induces bias in genome-wide studies and downstream analyses. In observational studies using UKBB data, participation bias has already been shown to distort phenotypic exposure-outcome associations^11,12,14^. If study participation includes a genetic component, biased estimates are also expected in genetic studies^15^. In gene-discovery studies, non-random participation may distort the association between a genetic variant and the outcome (cf. Fig 1A). In Mendelian Randomization, participation bias could induce an association between genetic instruments and unmeasured confounders of the exposure-outcome association, thereby violating a key assumption of the method (cf. Fig 1B/C). Recent genome-wide studies looking at proxies of participation bias have already described a genetic component to participatory behaviour and questionnaire responding^16–23^, implicating that genetic studies are not immune to bias. While much of the recent GWA output is produced by non-representative biobanks (e.g., UKBB, Million Veteran Program, 23andMe), the extent to which gene-discovery and downstream analyses are subject to participation bias is currently unknown.

**Figure 1.**
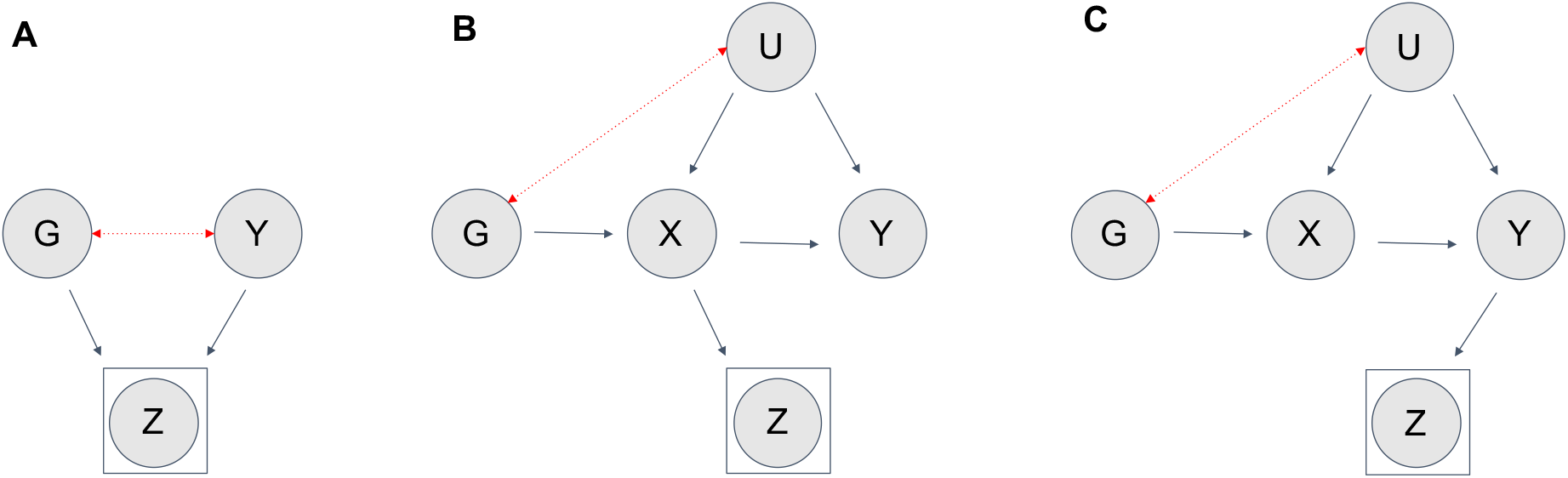
Illustration of the impact of participation bias in genetic studies Illustration of the relationships between a genetic variant (G) and an exposure (X) or outcome (Y) and study participation (Z). Panel (**A**) illustrates the effect of participation bias in genome-wide association studies, where (Z) is a common consequence of (G) and (Y) (dashed red). Conditioning on a common consequence (Z) induces a non-causal association between the genetic variant (G) and the outcome (Y). Panel (**B/C**) illustrates the effect of participation bias in Mendelian Randomization studies, where bias occurs if (Z) is a consequence of either the exposure (X) (Panel **B**) or the outcome (Y) (Panel **C**). Conditioning on (Z) induces an association between the genetic variant and confounders, thereby violating the MR assumption of exchangeability.

Participation bias is eliminated by the use of samples that are representative of their target population. To achieve representativeness in the UKBB, we derive a model for participation probability and create a pseudo-sample of the UKBB matching its target population. Thereby, it is possible to evaluate how a shift towards representativeness impacts genome-wide findings and downstream analyses. We anticipate that these findings will help to characterize the impact of participation bias in large volunteer-based samples used for biomedical research and to pin down areas of research that might be particularly susceptible to bias when relying on non-representative samples.

## Methods

First, we derived a model for participation probability by comparing 14 harmonized characteristics of the UKBB sample to that of a representative sample. Utilising the estimated participation probabilities, we conducted inverse probability weighted genome-wide association analyses (wGWA) on 19 UKBB traits. Second, to explore the genetic basis of UKBB participation, we conducted a GWA on the participation probability and evaluated the genetic findings. Finally, comparing wGWA results to those obtained from standard GWA analyses, we assessed the impact of participation bias on the estimation of three frequently studied quantities: 1) effect of genetic markers on complex traits, 2) heritability and genetic correlation estimates and 3) exposure-outcome associations obtained from Mendelian Randomization

## Samples

### UK Biobank

The UK Biobank (UKBB) is a large-scale prospective population-based research resource focusing on the role of genetic, environmental and lifestyle factors in health outcomes in middle age and later life. More than 9,000,000 men and women between 40 and 69 registered with the UK NHS were invited to take part. Of those, 5.4% (~500,000 individuals) were recruited in 22 assessment centres across England, Wales and Scotland between 2006 and 2010^24,25^. Included in this study were data from UK Biobank participants of European ancestry passing standard GWA analysis quality control measures^26^. We further filtered the sample according to geographical region (excluding individuals from Scotland and Wales) to match the geographical regions included in the reference sample (HSE), and removed individuals with missing data in auxiliary variables used to generate the propensity scores (further described below).

### Health Survey England

The Health Survey for England is an annual probability sample set out to measure health and related behaviours in a nationally representative sample of adults and children living in private households in England^27^. In our study, we included data from five cohorts recruiting a sample of more than 80,000 individuals between 2006 and 2010 (i.e., the UKBB recruitment period). We applied the same inclusion criteria to HSE data as used for UKBB recruitment, retaining only individuals aged between 40 and 69 years and of (self-reported) European ancestry. HSE response rates ranged between 64% and 68%^27^. HSE sample weights are supplied to account for the unequal probabilities of selection and non-response^28^, weighing individuals as a function of sex, household type, region and social class. In this study, the HSE weights were incorporated in LASSO regression predicting UKBB participation (described below).

### UK Census data

We also exploited data from the 2011 Census Microdata, a 5% sample of anonymized individual-level Census record^29^, which runs every ten years to collect basic demographic variables (e.g., educational attainment, age, general health) through a paper-based or online questionnaire. With a 95% response rate, the UK Census microdata is highly representative of the UK population. We applied the same selection criteria to the Census data as to the UKBB and HSE (i.e., filtered according to geographical region, restriction of individuals to European ancestry and ages 40-69), resulting in a relevant sample of n=895,649. We extracted all variables that could be harmonized with the UKBB and HSE data (further described in Supplement). The Census data was solely used to assess the level of representativeness of the HSE, by comparing the distributions and associations between variables present in both the HSE and Census sample. For the generation of UKBB probability weights, we rather used the HSE sample, given its richer phenotypic data critical for accurate weight estimation.

## Analysis

### Auxiliary variables

We adjust for participation bias in the UKBB using probability weighting^30^. This approach adjusts for non-response bias by weighting over-and under-represented individuals, thereby creating a pseudo-population that is representative of its target population^31^. Probability weighting relies on auxiliary variables available for both a selected (non-representative) and a representative reference sample. In this study, we selected auxiliary variables tapping into dimensions related to health, lifestyles, education and basic demographics. We included all variables that could be harmonized across the two datasets (HSE and UKBB) with few missing observations (i.e., < 50,000 in the UKBB, < 500 in HSE). Fourteen variables derived from twelve measures were included and harmonized across the two datasets. Five continuous variables included: age, BMI, weight, height and education (age when completed full-time education). Nine categorical variables included: household size (1, 2, 3, 4, 5, 6 or 7 or more), sex (male/female), alcohol consumption frequency (never/few times per year/monthly/once or twice weekly/three or four times weekly/daily), smoking status (never/previous/current), employment status (employed/economically inactive/retired/unemployed), income (<18k/18k−31k/31k−52k/52k−100k/>100k), obesity status (underweight/healthy weight/overweight/obese) and overall health (poor/fair/good). Further details of the coding of the variables in each dataset are provided in the Supplement.

### Construction and evaluation of UK Biobank probability weights

To derive the model for participation probability, we first combined the harmonized UKBB data with the data from the reference sample (HSE). We then used LASSO regression in glmnet^32^ to predict UKBB participation (*P*_*i*_, with UKBB = 1; HSE = 0), conditional on the harmonized auxiliary variables described above. We included fourteen main effects (five continuous variables, nine binary/categorical variables) in the model. All categorical and binary variables were entered as dummy variables, indexing each possible level of the variable. In addition, we included all possible two-way interaction terms among the dummy and continuous variables, resulting in 903 included predictors. LASSO performs variable selection by shrinking the coefficients for variables that contribute least to prediction accuracy. The shrinkage is controlled by the tuning parameter (*λ*), which was obtained using 5-fold cross-validation that minimizes the cross-validated error.

The predicted probabilities (P_i_) were then used to build the individual sampling weights (*w*_*i*_). The weights were calculated as an extension of standard Inverse Probability Weights [*w*_*i*_ = (1-P_i_)/P_i_], designed to make the weighted sample estimates conform to the population estimates^31^. To assess the performance of the generated weights, we evaluated the extent to which the weighting recovered means (for continuous variables) and prevalences (for binary traits) in the UKBB and, hence, mitigated participation bias. We also quantified participation bias as the differences between the correlations among all auxiliary variables within the UKBB (*r*_UKBB_) and the HSE (*r*_HSE_). The degree to which the weighted correlations (*r*_UKBB_W_) reduced bias was estimated as (|*r*_HSE_ -*r*_UKBB_| -|*r*_HSE_ -*r*_UKBB_W_|) / (|*r*_HSE_ - *r*_UKBB_|), where a value of one indicates that weighting fully eliminated bias. The weighted means (and proportions) for a given variable (*X*_*i*_) were estimated using the weights (*w*_*i*_), with the expression: 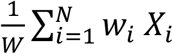 where 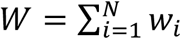.

We further evaluated if over-fitting was a problem, by re-running LASSO in train-test splits of the data (5-fold leave-one-out cross-validation, with a split ratio of 80:20). Here, we used the training sample (80% of the data) for model estimation and the test sample (20% of the data) to generate the out-of-sample predicted probabilities. The degree of participation bias reduction was then compared between the out-of-sample predicted probabilities and the full sample probabilities.

### Probability weighted genome-wide association analyses

To evaluate the extent to which SNP effects were distorted by participation bias in the UKBB, we conducted inverse probability weighted genome-wide association analyses (wGWA). wGWA was performed for 19 UKBB health-related traits collected at baseline with few missing observations (n_missing_<50,000). The coding of all variables, genotyping, imputation and quality control (QC) procedures are described in the Supplement. Additional QC filters for genome-wide analyses were applied to select participants (i.e., exclusion of related individuals, exclusion of non-White British ancestry based on principal components, high missing rate and high heterozygosity on autosomes) and genetic variants (Hardy–Weinberg disequilibrium *P* > 1 × 10^−6^, minor allele frequency >1% and call rate >90%).

We obtained unweighted SNP estimates 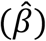 from a standard ordinary least squares (OLS) linear regression model. The weighted SNP estimate 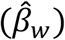 were obtained from weighted least squares (WLS) regression. All GWA analyses were conducted in LDAK (version 5.2)^33,34^, which was extended to accommodate sampling weights in a linear WLS model (--linear -- sample-weights). The standard least squares estimate of the variance is based on the assumption of homoskedasticity (i.e., that the residual variance is constant across individuals). Since the use of sampling weights violates this assumption, we used the Huber-White estimator^35^ to estimate the variance of the coefficients:

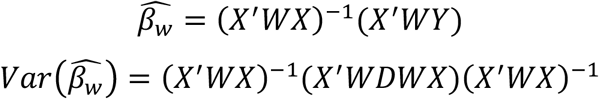

with

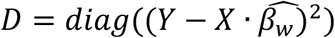

where *Y* denotes the phenotypic outcome vector, *W* is a diagonal matrix with the probability weights sitting on the diagonal and *X* is a column vector of the genotype values.

Both models included the same covariates (PC1-PC5, sex, age, batch effect). We applied a linear model to all outcomes (continuous and binary traits). This was done to allow for standardization of SNP estimates and to ensure comparability of effect sizes.

Two additional sets of analyses were conducted to explore the genetic basis of UKBB participation: First, we conducted autosomal wGWA and standard GWA on biological sex and evaluated if wGWA reduced sex-differential participation bias. As previously suggested^22^, autosomal heritability linked to biological sex could result from sex-differential participation. As such, reduced heritability estimates in wGWA compared to GWA would provide evidence for the utility of wGWA for participation bias correction. In addition, we compared the resulting SNP effects to the effects of previously identified sex-associated variants (*p*< 5 × 10^−8^). Here, 49 variants assessed in an independent sample of >2,400,000 volunteers curated by 23andMe^22^ were selected.

Second, we conducted a genome-wide analysis on the liability to UKBB participation, by including the individual participation probabilities as the outcome of interest in wGWA. Application of standard GWA analysis is not possible in this context, as this approach stratifies for the outcome of interest by selecting a subset of the population willing to participate. LD-independent SNPs reaching genome-wide significance (*p*<5×10^−8^) were selected via clumping (--clump-kb 250 –clump-r2 0.1). PhenoScanner^36^, a database of genotype-phenotype associations from existing GWA studies, was used to explore previously identified associations of lead SNPs with other phenotypes. Genetic correlations with other traits were estimated using LD-score regression^37^ as implemented in the R-Package GenomicSEM^38^. The summary statistic files used in LD-score regression were obtained for 49 health and behavioural phenotypes, using publically available summary statistic files accessible via consortia websites or the MRC-IEU OpenGWAS project (**Error! Hyperlink reference not valid**. (sTable 1 for details).

### LD score regression and heritability estimates

SNP heritability estimates were obtained for both the standard GWA and wGWA output (*h*^2^ and 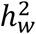, respectively), using LD score regression as implemented in GenomicSEM. We applied the default settings (restricted SNPs to MAF>0.01, LD-scores from the European-ancestry sample in the 1000 Genomes Project^40^). For binary phenotypes, the observed scale was converted to the liability scale^41^, where the population prevalence was set to be equal to the weighted prevalence in the UKBB. We also estimated bivariate genetic correlations among all phenotypes included in standard GWA and wGWA (*rg* and *rg* _*w*_, respectively). To compare the estimates obtained from wGWA and standard GWA, we calculated the difference (*rg* _*DIFF*,_ = *rg*– *rg* _*w*_ and 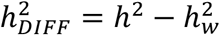) and used the following test statistic (here exemplified for *rg* _*DIFF*_):

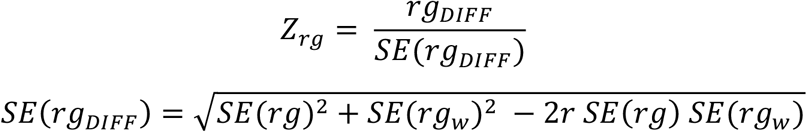

The correlation coefficients 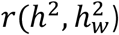 and *r*(*rg, rg* _*w*_) were obtained from 200-block Jackknife analysis. For this, we split the genome into 200 equal blocks of SNPs and removed one block at a time to perform Jackknife estimation.

### Mendelian Randomization analyses

To evaluate the impact of selection bias when using MR, we assessed if sample weighting altered MR estimates. As genetic instruments, we selected LD-independent (--clump-kb 10,000 --clump-r2 0.001) SNPs reaching genome-wide significance (*p*<5×10^−8^) in either wGWA or standard GWA for a given phenotype. Phenotypes with few (<10) genetic instruments were not included in MR analyses. We used the inverse-variance weighted (IVW) MR estimator, which combines the ratio estimates of the individual genetic variants *G*_*j*_ to derive the causal effect 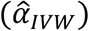. The ratio estimate is 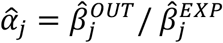, where 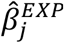 corresponds to the SNP-exposure association and 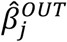 corresponds to the SNP-outcome association. Since the IVW estimator assumes that the uncertainty in the genetic association with the exposure is zero, we used the following correction^42^ to account for selected genetic variants 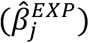 that were genome-wide significant in one analysis (e.g., standard GWA) but not the other (e.g., wGWA) for the same trait: 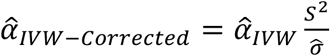 where 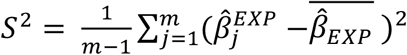, and 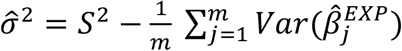 where *m* refers to the number of SNPs selected as instruments. The corresponding variance was estimated as: 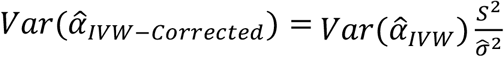.

For each exposure-outcome association, we obtained 1) an MR estimate using the SNP effects from standard GWA analyses and 2) an MR estimate using the SNP effects from wGWA analyses. We included in MR the standardized SNP effects and standard errors (i.e., effect of the genotype on the standardized outcome), which were derived using the following formula^43^: 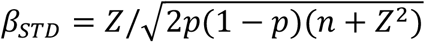 and 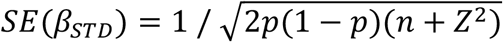, where *n* is the sample size, *p* is the minor allele frequency (MAF), and *Z* is the SNP effect 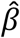 divided by its standard error 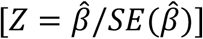. Of note, when standardizing the weighted estimates 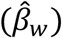, *n* was replaced by the effective sample size 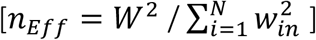 to account for the unequal contribution per observation. *w*_*in*_ refers to the normalized probability weights, obtained by dividing *w*_*i*_ by its mean 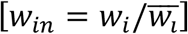.

To compare the standard 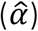 to the weighted MR 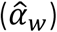 estimates, we estimated 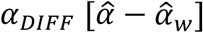 and the corresponding test statistic as *Z* = *α*_*DIFF*_ / *SE*(*α*_*DIFF*_), where 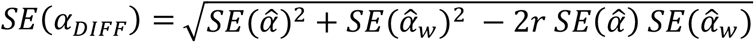.The correlation coefficient was derived using a Jackknife procedure, where we performed MR leaving out each SNP in turn to then calculate the correlation 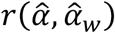 from these results. The results were corrected for multiple testing using FDR-correction (controlled at 5%), correcting for the total number of conducted MR analyses.

## Results

### Samples

From the five HSE cohorts comprising a total sample of n=81,118, we retained n=22,646 after applying the same inclusion criteria used for UKBB recruitment. After further exclusion of HSE individuals with missing data on the 14 auxiliary variables, we included a final sample of n=21,816. Comparing the distribution of a subset of auxiliary variables also available in the UK Census Microdata (n=895,649) shows that the profile of the HSE sample closely matches that of the Census sample (sTable 2). More specifically, proportions were comparable between HSE and Census, but deviated in the UKBB for most of the selected variables, such as female gender (P_CENSUS_=51%, P_HSE_=51%, P_UKBB_=54%), individuals of age ≥65 (P_CENSUS_=13%, P_HSE_=13%, P_UKBB_=19%), mean age when completed full time education (M_CENSUS_=16.6, M_HSE_=16.4%, P_UKBB_=17.2%), retired individuals (P_CENSUS_=19%, P_HSE_=19%, P_UKBB_=34%). Further inspection of the associations between variables available in the HSE and UK Census (sFigure 1) highlights that the HSE captures well the characteristics of the population residing in England.

Of the initial UKBB sample (502,645 participants), we excluded individuals of age >69 and <40 (n=2463), individuals from Scotland or Wales (n=56,483), individuals of non-European ancestry (n=28,371), individuals withdrawing consent (n=161) and individuals with missing data on any of the auxiliary variables (n=21,868). The sampling weights were generated for n=393,299 UKBB individuals, of which 109,550 were removed after applying QC steps for genome-wide analyses.

### Performance of the UK Biobank probability weights

Figure 2A shows the distribution of the normalized propensity weights (*w*_*in*_) for UKBB individuals. The probabilities used to construct the weights were obtained from a LASSO regression model retaining 454 of the 903 initially included predictors. Figure 2B illustrates which auxiliary variables most strongly linked to UKBB participation (UKBB = 1; HSE = 0), highlighting that older (retired) and more educated non-smoking individuals were particularly likely to participate.

**Figure 2.**
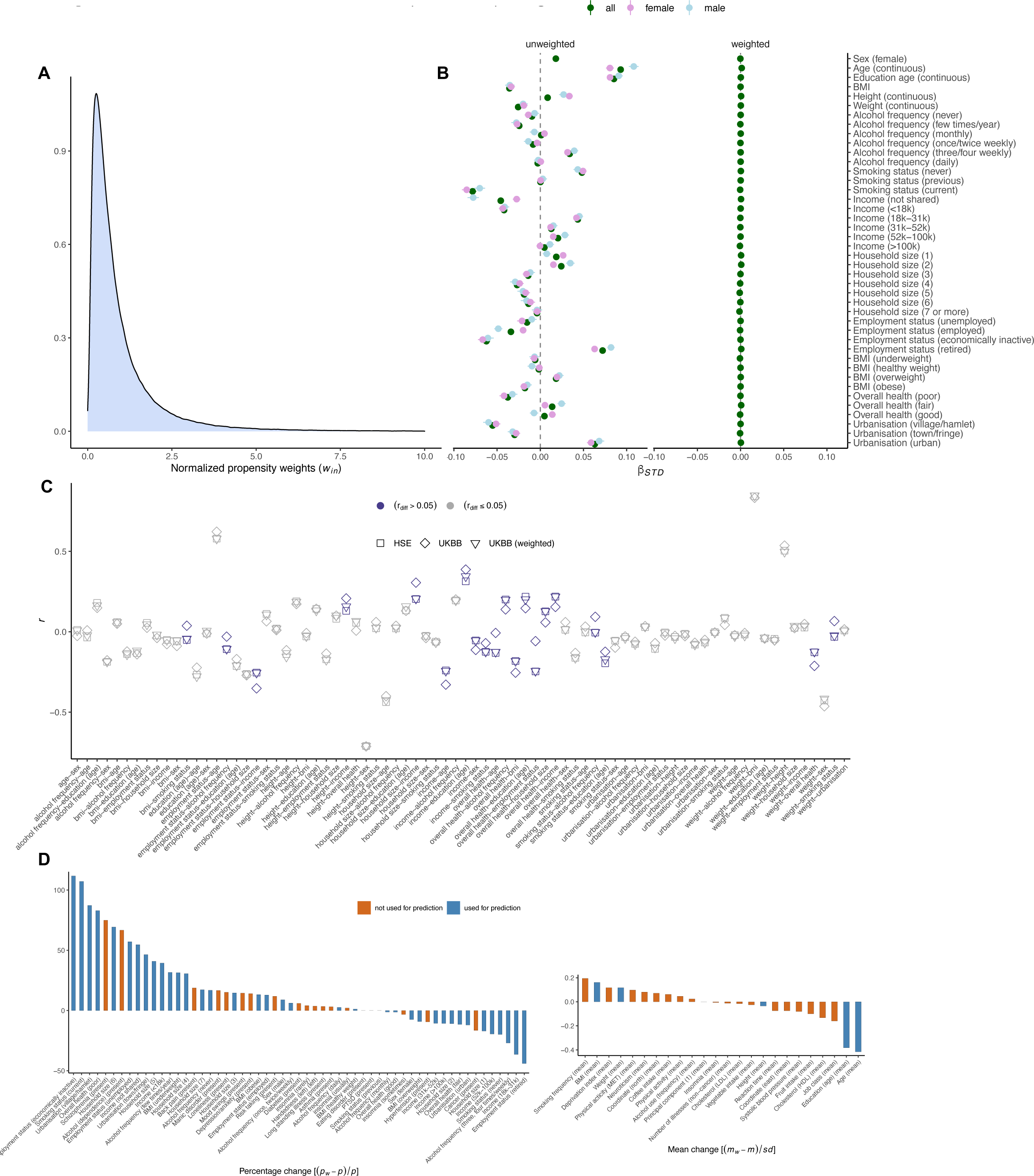
Performance of the UK Biobank probability weights Panel (**A**) presents the truncated density curves of the normalized propensity weights (*w*_*in*_) for UKBB participants, ranging from 0.02 to 50.01. Panel (**B**) shows standardized coefficients of variables predicting UKBB participation (HSE = 0; UKBB = 1) in univariate logistic regression models. Coefficients are provided for all UKBB participants and males and females separately. Panel (**C**) plots the correlation coefficients among all auxiliary variables within the UKBB (obtained from weighted and unweighted analyses), and within the HSE. Highlighted in blue are results where the coefficients between the UKBB (*r*_UKBB_) and the reference sample (*r*_HSE_) deviated (*r*_diff_ >0.05, where *r*_diff_=|*r*_HSE_-*r*_UKBB_|). Panel (**D**) depicts the percentage change (for categorical variables) and change in means as a function of weighting, obtained for a number of health-related UKBB phenotypes, including the auxiliary variables (blue) and variables not used to construct the weights. Percentage change was estimated as the difference between the weighted (*p*_*w*_) and unweighted proportion (*p*), divided by the unweighted value [(*p*_*w*_ – *p*) / *p* × 100]. Change in means was expressed as a standardized mean difference, estimated as the difference between the unweighted mean (*m*) and the weighted mean (*m*_*w*_), divided by the unweighted standard deviation (sd) [*m*_*w*_-*m*/*sd*].

To evaluate the performance of the weights, we first assessed if probability weighting recovered the reference (HSE) population distributions. We included the generated weights in univariate logistic regression model predicting UKBB participation, where UKBB individuals were given their normalized weight (*w*_*in*_) and HSE participants were given a weight of 1. When applying probability weighting (shown on the right side of Figure 2B), previously significant predictors became non-significant. All means and proportions in the HSE, UKBB (unweighted) and UKBB (weighted) are provided in sTable 3. Next, we estimated the degree of bias reduction following probability weighting. Here, we quantified participation bias as the difference between an estimate of association obtained in the UKBB (*r*_UKBB_) and the reference sample (*r*_HSE_). The largest difference [*r*_diff_=|*r*_HSE_-*r*_UKBB_|] was present for employment status with overall health (*r*_diff_=0.19; *r*_HSE_=-0.25; *r*_UKBB_=-0.06), overall health with age (*r*_diff_=0.12; *r*_HSE_=-0.13; *r*_UKBB_=-0.01), household size with income (*r*_diff_=0.10; *r*_HSE_=0.20; *r*_UKBB_=0.31), and employment status with income (*r*_diff_=0.10; *r*_HSE_=-0.25; *r*_UKBB_=-0.35) (cf. Figure 2C). Application of probability weighting eliminated most bias induced by selective participation (median bias reduction: 0.97; mean: 0.91, range: 0.58 - 0.998). The estimates were very similar to the cross-validated model (median bias reduction: 0.96; mean: 0.90, range 0.50; 0.998), highlighting that over-fitting was unlikely to be a problem.

Finally, Figure 2D summarizes changes in means and proportions as a result of probability weighting, estimated for the auxiliary variables (in blue), as well as other UKBB variables (in orange) not used to construct the weights. Application of weighting resulted in overall more unfavourable health outcomes and demographics, including increases in mental illness (higher rates of schizophrenia and alcohol addiction) and poorer socioeconomic status (higher deprivation index, lower job class), providing further indication that weighting largely reduced the healthy volunteer bias of the UKBB.

In summary, probability weighting successfully created a pseudo-sample of the UKBB achieving higher levels of representativeness, thus providing a useful tool for participation bias correction in downstream analyses.

### Probability weighted genome-wide association analyses on UK Biobank traits

The effect of participation bias on genome-wide results was evaluated by comparing the SNP estimates obtained from probability weighted genome-wide analyses, wGWA (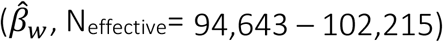, N_effective_= 94,643 – 102,215), to standard GWA analyses 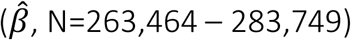, N=263,464 – 283,749). As illustrated in Figure 3, the impact of participation bias was assessed in terms of changes in effect sizes across SNPs and genome-wide discovery (i.e., number of identified SNPs).

First, Figure 3A highlights the number of SNP where bias induced overestimation 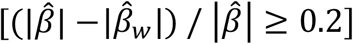 or underestimation 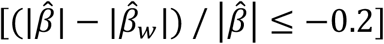 of SNP effects. Among all genome-wide hits (1690, with *p*< 5 × 10^−8^), overestimation was more common (420 SNPs, 24.85% of all genome-wide SNPs) than underestimation (290 SNPs, 17.16% of SNPs). More specifically, underestimation was most common for cancer (57% of SNPs), loneliness (50%), education (33%) and reaction time (33%), whereas overestimation was present for depression/anxiety (67%), coffee intake (63%) and smoking status (58% of SNPs). There was no evidence of change in direction of effects as a result of participation bias (cf. sResults, Supplement). Second, with respect to genome-wide discovery (Figure 3B), we found that of all SNPs identified either in wGWA or GWA analyses (n=1690 across all phenotypes), 25 SNPs (1.48%) were missed as a result of participation bias, as these SNPs reached significance only in the weighted analyses. Novel SNPs were identified for 12 of the 19 included traits, most notably for depression/anxiety (50% of genome-wide SNPs linked to depression/anxiety were missed by standard GWA), cancer (29%) and loneliness (25%). Detailed results are listed in sTable 4 and plotted in sFigure 2-3.

**Figure 3.**
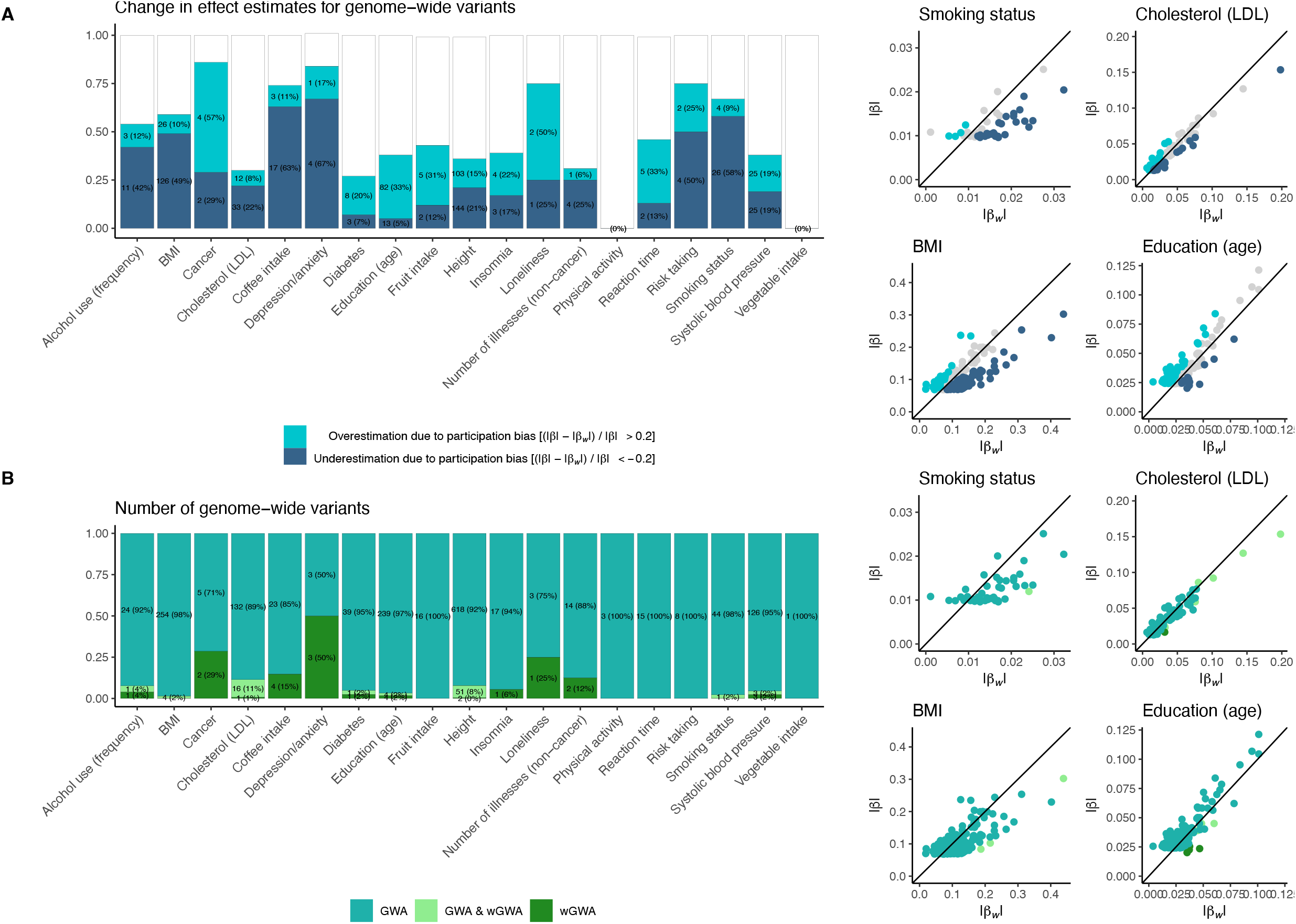
SNP estimates from weighted and unweighted genome-wide analyses Summary of comparison between SNP effects obtained from probability weighted genome-wide association (wGWA) and standard GWA analyses on 19 traits. Panel (**A**) summarizes the proportions of overestimated and underestimated SNP effects as a result of participation bias. Shown in panel (**B**) are the numbers and proportions of SNPs reaching genome-wide significance in either standard GWA, wGWA or both (GWA & wGWA). The scatter plots to the right plot the weighted (|*β*_*w*_|) against the unweighted (|*β*|) SNP effects for four selected traits.

### Probability weighted genome-wide association analysis on sex

UKBB participants were proportionately more female (female_UKBB_=54.38%) compared to the general population (female_HSE_=50.74%; female_CENSUS_=50.62%). Probability weighting recovered the population prevalence in the UKBB (weighted female_UKBB_=50.36%). SNP heritability estimates (*h*^2^) (sFigure4A) using wGWA led to almost half of that of the standard GWA (*h*^2^ on liability scale = 1.2%, *p*=0.1 in wGWA versus 2.1%, *p*=5.4e-11 in standard GWA). sFigure 4B/sTable 5 display the SNP effects of 49 variants previously associated with sex

(*p*< 5 × 10^−8^, in an independent sample of >2,400,000 volunteers) to estimates obtained from standard GWA and wGWA. Of those, 18 (36.73%) SNPs showed significantly reduced sex-associated effects in wGWA. In contrast, only 3 (6.12%) SNPs showed reduced sex-associated effects in standard GWA.

### Genome-wide association study on the liability to UKBB participation

wGWA on UKBB participation was conducted in N_eff_=102,215 participants. 28 SNPs reached genome-wide significance (*p*< 5 × 10^−8^), of which LD-independent 23 SNPs were selected after clumping. Figure 4A shows the Manhattan plot with positional mapping of genome-wide SNPs associated with the liability to UKBB participation (cf. sTable 6 for annotation and estimates of significant SNPs). The QQ plot (sFigure 5) can be found in the Supplement. SNP heritability for UKBB participation was *h*^2^=0.009 (se=0.005) (LD-score intercept: 1.055). A lookup of SNP-trait associations estimated in previous GWA analyses showed that UKBB participation-associated variants mostly tapped into age-related outcomes (e.g., cause of death: cancer/dementia/fatty liver disease/pneumonia) (sTable 7). LD score regression analyses (cf. Figure 4B, sTable 8) implicated substantial genetic correlations between UKBB participation and phenotypes related to socioeconomic factors and previously assessed participatory behaviour, including educational attainment (rg=0.92), income (rg=0.81), participation (response to email invitationand mental health survey completion) (*rg*=0.75 and *rg*=0.7, respectively), intelligence (*rg*=0.73) and cigarette use (age of onset) (*rg*=-0.72).

**Figure 4.**
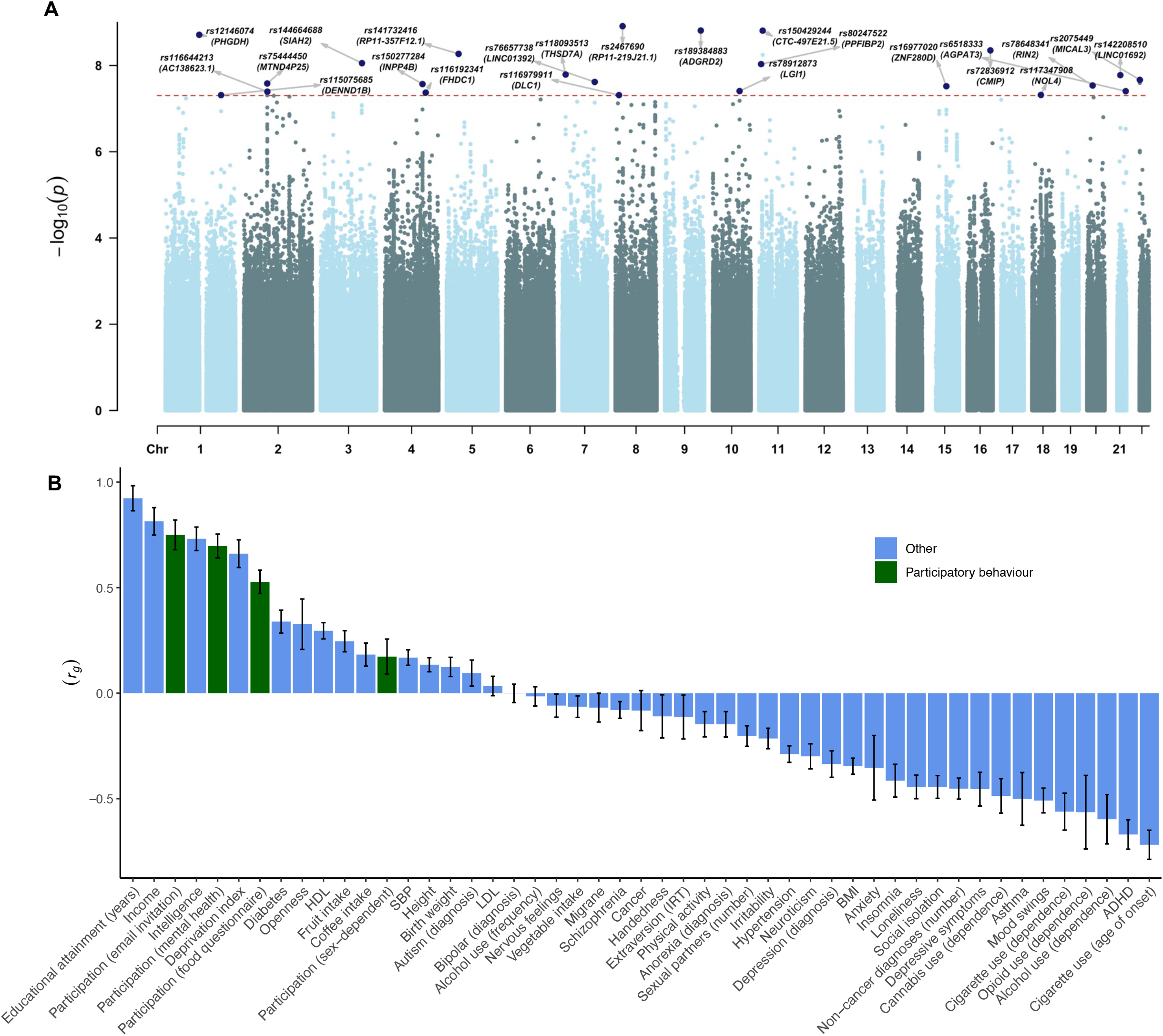
Genome-wide association study on the liability to UK Biobank participation Panel (**A**) displays the Manhattan plot of the genome-wide association study on the liability to UKBB participation. Labels are provided for the top LD-independent genome-wide significant SNPs (i.e., SNPs above the horizontal line, with *p*<5×10^−8^) and gene names obtained through positional mapping. The x-axis refers to chromosomal position, the y-axis refers to the *p*-value on a −log10 scale. Panel (**B**) shown are the genetic correlations (*r*_*g*_) of the UKBB participation with traits indexing participatory behaviour (in green) and other traits (in blue).

### Weighted SNP heritability and genetic correlation estimates

Bias was assessed in terms of differences in SNP heritability 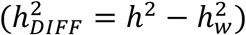 and genetic correlations (*rg*_*DIFF*_ = *rg* − *rg*_*w*_) between standard and weighted GWA analyses (Figure 5). On average, heritability estimates differed by 1.5% (liability scale 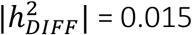, range 0 - 0.05). 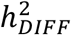 was highest for BMI 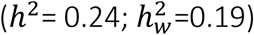, education (*p*_FDR_<0.05) 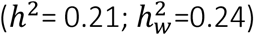 and diabetes 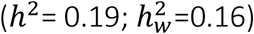. Of all assessed traits included in LDSC regression (n=18), five showed significant (*p*_FDR_<0.0.5) 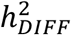, of which four (80%) were over-estimated and one (education) was under-estimated as a result of participation bias. The weighted and unweighted heritability estimates are plotted in sFigure 6 and additional statistics (e.g., LDSC intercepts) are provided in sTable 9.

**Figure 5.**
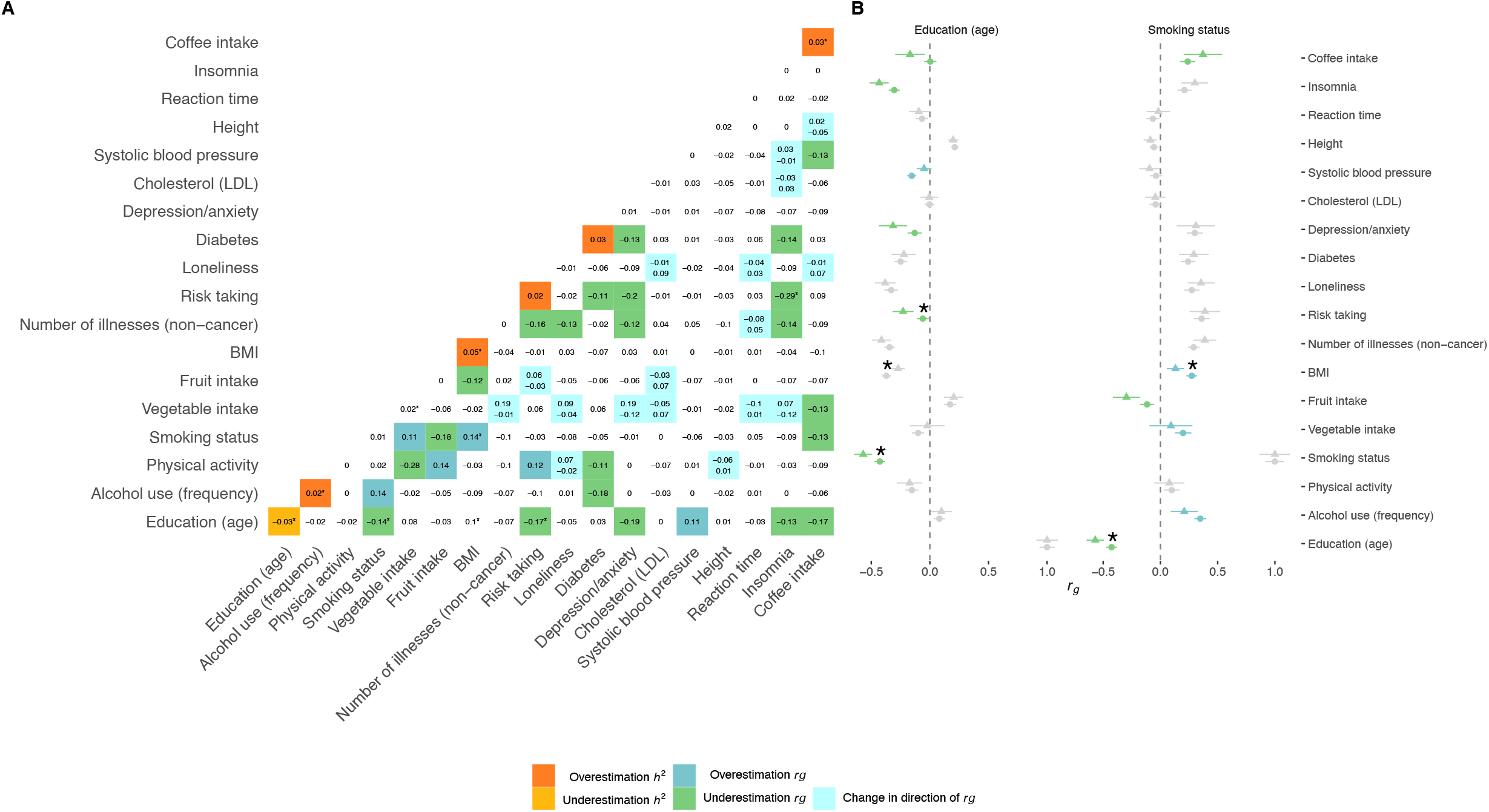
Weighted SNP heritability and genetic correlation estimates Plotted in Panel (**A**) are the differences in SNP heritability (*h*^*2*^_*DIFF*_ = *h*^*2*^ - *h*_*w*_^*2*^) and genetic correlations (*rg*_*DIFF*_ = |*rg*| *-*|*rg*_*w*_|) obtained from weighted and standard GWA analyses. The diagonal shows the differences in SNP heritability, where bias leading to overestimation (*h*^*2*^_*diff*_ > 0.02) are plotted in orange and bias leading to underestimation (*h*^*2*^_*diff*_ < −0.02) are plotted in yellow. The off-diagonal highlights overestimated genetic correlations (*rg*_*DIFF*_ > 0.1) in blue and underestimated genetic correlations (*rg*_*DIFF*_ < −0.1) green. Tiles coloured in turquoise index genetic correlations where *rg* and *rg*_*w*_ show opposite directions (with *rg* printed at the top and *rg*_*w*_ printed at the bottom of the tile). Panel **B** illustrates estimates of genetic correlations (*rg* shown as ⬤; *rg*_*w*_ shown as▲) and the corresponding confidence intervals for two selected traits. (*) Estimates showing significant differences (*p*_FDR_<0.05).

Concerning estimates of genetic correlations, we found an average difference of |*rg*_*DIFF*_| = 0.07 (range 0 – 0.31) between results obtained from standard and weighted GWA analyses. Participation bias leading to *rg-*overestimation was most notable for *rg*(BMI, smoking status) [*rg*=0.27; *rg*_*w*_=0.13], *rg*(fruit intake, physical activity) [*rg*=0.32; *rg*_*w*_=0.18] and *rg*(alcohol use frequency, smoking status) [*rg*=0.35; *rg*_*w*_=0.21]. *rg-*underestimation was most prominent for *rg*(insomnia, risk taking) [*rg*=0.02; *rg*_*w*_=0.31], *rg*(vegetable intake, physical activity) [*rg*=0.3; *rg*_*w*_=0.58] and *rg*(depression/anxiety, risk taking) [*rg*=0.27; *rg*_*w*_=0.47]. For five (3.27%) of the assessed trait pairs (n=153) the weighted and standard genetic correlations were significantly (*p*_FDR_<0.05) different, of which education was the most commonly implicated trait (sFigure 7, sTable 10). Change in direction of genetic correlations as a result of participation bias was less present (cf. sResults, Supplement).

### Effect of participation bias on Mendelian Randomization estimates

Figure 6 summarizes which MR estimates were affected by participation bias, as inferred from differences between the standard and weighted MR estimates 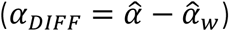. On average, participation bias led to an absolute change of 0.038 in standardized MR estimates (range 0 – 0.15). Most affected were associations between lifestyle choices, including coffee intake on BMI 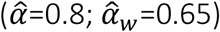, fruit consumption on LDL cholesterol 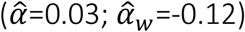 and fruit consumption on coffee intake 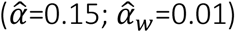 (sFigure8 and sTable 11). Of all exposure-outcome associations tested (k=234), 14 (6%) estimates were either overestimated (2%, 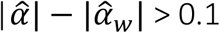) or underestimated (4%, 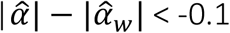). Significant (*p*_FDR_<0.05) differential effects were only present for two of the exposure-outcome associations tested (education on BMI; smoking status on fruit consumption). There was little evidence of bias resulting in changes in direction of MR estimates (sResults, Supplement).

**Figure 6.**
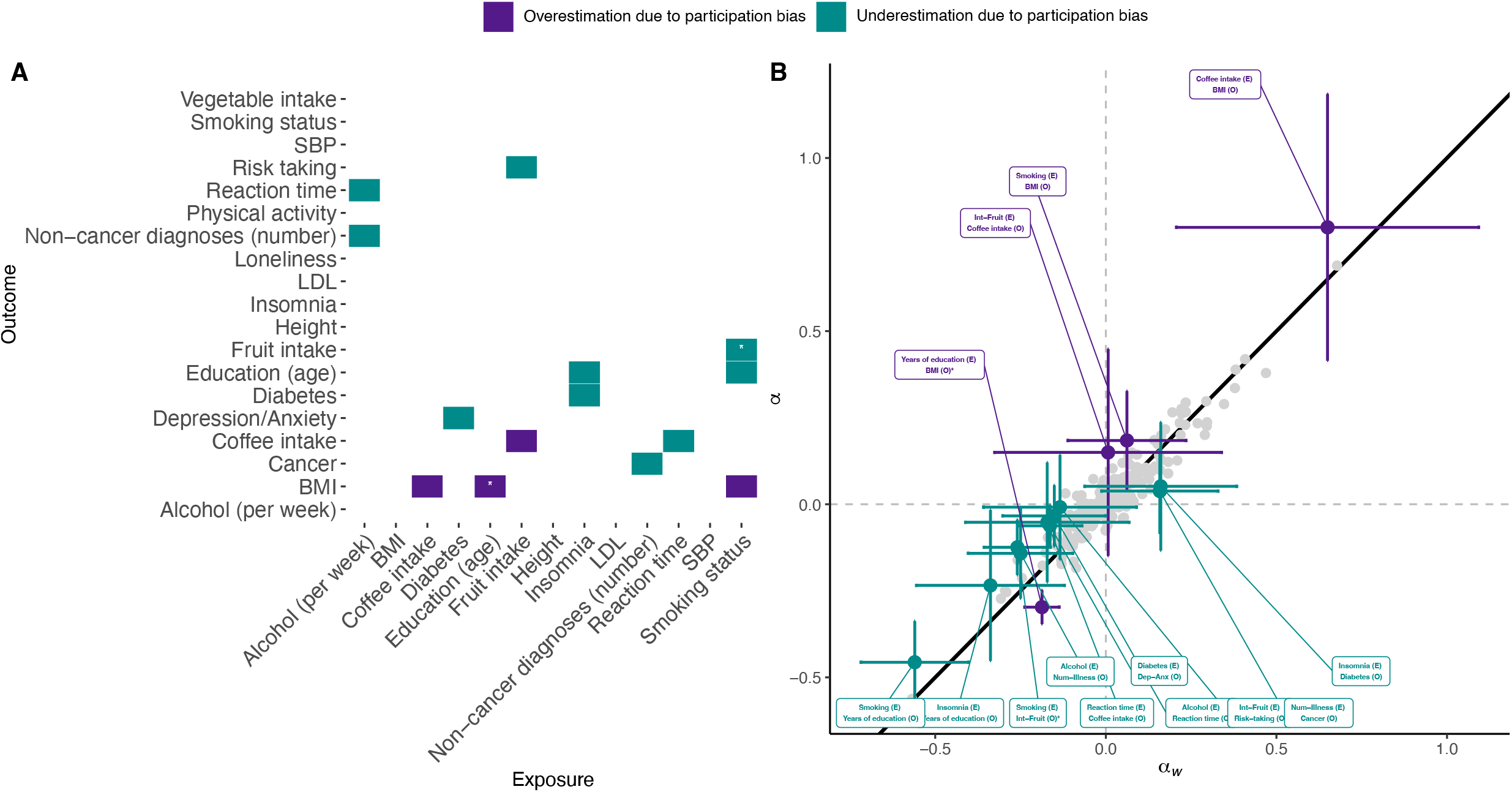
Effect of participation bias on MR estimates of exposure-outcome associations Summary of results obtained from weighted 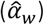 and standard 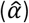 Mendelian Randomization (MR). MR estimates subject to overestimation 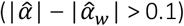 as a results of participation bias are highlighted in violet. MR estimates subject to underestimation 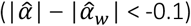 are highlighted in cyan. The asterisks (*) highlight results where 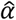 and 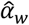 showed significant (*p*_FDR_<0.05) differences.

## Discussion

While the use of large volunteer-based biobanks is key to advancing genetic epidemiology, it is currently unclear to which extent selective participation impacts genotype-phenotype association obtained from the data. In this work, we conducted inverse probability weighted GWA (wGWA) on 19 traits to correct for participation bias in genome-wide analyses. We found that participation bias can distort genome-wide findings and downstream analyses.

Overall, the bias mostly affected magnitude of effects, rather than direction, but was present across all sets of genome-wide analyses. For genome-wide association analysis, we find that estimates are subject to both overestimation (e.g., for cancer and education) and underestimation (e.g., for coffee intake, depression/anxiety). wGWA also led to the discovery of novel loci for 12 of the included traits, highlighting SNPs that would be missed as a result of participation bias. Of note, although effect size estimates can shrink as a consequence of participation bias, the increased standard errors of the inverse probability weighting (due to reduced effective sample size) renders new discoveries difficult. Participation bias also distorted heritability estimates, genetic correlations and Mendelian Randomization estimates, most notably for socio-behavioural traits including education, diet, smoking and BMI. In contrast, more molecular/physical traits (e.g., LDL, SBP) exhibited less bias as a result of selective participation. Such pattern is in line with existing studies^22,23^ and our findings of high genetic correlations between the liability to UKBB participation and socio-behavioural traits, particularly education, income and substance use. More broadly, *different* sources of bias are likely to affect *similar* phenotypes in genome-wide studies, in that socio-behavioural phenotypes are subject to bias resulting from selective participation^22,23^, indirect genetic effects^3^, assortative mating^4^, error in measurements^44^ and population stratification^45^.

Our work builds and extends recent efforts evaluating bias due to selective participation. More specifically, we replicate findings showing that phenotypic exposure-outcome associations in the UKBB differ from those estimated in probability samples^12,14^: participation bias, defined as the difference in exposure-outcome associations in the UKBB and the reference sample (HSE), was substantial for a number of associations. For example, phenotypically, the impact of participation bias was most prominent for the link between overall health with age and employment status. Application of probability weighting eliminated a significant proportion (>90%) of bias due to selective participation in the UKBB. To our knowledge, this work is the first to provide participation bias corrected GWA on complex traits related to health and behaviour. We highlight patterns of bias and point to areas of research that are most impacted by this bias. Since genome-wide association summary statistics are increasingly used in epidemiological research to study causal questions concerning education, diet and behaviour, greater care should be taken when relying on data obtained from non-random samples. Biobank data in which participation bias cannot be assessed (e.g., in self-selection samples without a defined target population) may therefore be only of limited utility when scrutinizing genotype-phenotype relationships. As part of this work, we provide software to perform wGWA, which allows researchers to conduct sensitivity checks when relying on non-representative samples. Alternatively, recruitment schemes incorporating probability sampling can help reduce bias, but samples are typically small given the substantial costs associated with recruitment.

This study comes with a number of shortcomings. First, while the application of probability weighting successfully removed bias resulting from selective participation in the UKBB, residual bias may still exist. Important factors independently predicting UKBB participation may have been missed when modelling participation probability, which would reduce the efficacy of probability weighting. When choosing a reference population, there is a trade-off between representativeness of the reference sample and the number of available variables to match the samples. We choose to use the HSE as a reference sample to strike a balance between these two factors, but biases can remain if important variables were not measured or the reference sample is not representative enough. Second, genome-wide analyses were restricted to phenotypes with little missing data. This is a shortcoming since traits with substantial missing data are perfect candidates for characteristics influencing participation.

As such, we did not evaluate the impact of participation bias on variables collected at follow-up. Finally, the UKBB probability weights are sample specific, constructed for a more educated, healthy, older, and female sample residing in England. Bias due to selective participation will differ across study contexts and participation mechanisms as evaluated in this study are therefore not generalizable to other cohorts.

In conclusion, our results highlight that selective participation can lead to bias in genome-wide results and downstream analyses, most visibly for socio-behavioural traits. Moving forward, more efforts ensuring either sample representativeness or methods correcting for participation bias are paramount, especially when studying the genetic underpinnings of behaviour, lifestyles and educational outcomes.

## Data availability

Standard and probability weighted UK Biobank association statistics, computed using LDAK version 5.2, will be made available through the GWAS catalog.

## Code availability

LDAK is available at http://dougspeed.com/downloads/ All analytical scripts are available at https://github.com/TabeaSchoeler/TS2021_UKBBweighting

## Acknowledgements

This research has been conducted with the UK Biobank Resource under application number 16389; we thank all biobank participants for sharing their data. We are grateful to all participants involved in the Health Survey England and the 2011 Census Microdata, and we thank the Office for National Statistics for granting access to the data. This study would not have possible without the use of publicly available genome-wide summary data and software tools. The authors gratefully acknowledge these resources, and thank the research participants, the research teams and institutions that have contributed to this research.

Computations have been performed on the HPC cluster of the Lausanne University Hospital. The authors thank Yves Tillé for the helpful discussions and relevant comments.

