## Supplement for "Correction for participation bias in the UK Biobank reveals non-negligible impact on genetic associations and downstream analyses"

### Supplement Material


SMETHODS 2

Coding of variables included from the UKBB and HSE 2

Coding of variables harmonized across the UKBB, HSE and the UK Census Microdata 3

Genotyping, imputation and quality control in the UK Biobank 4

SRESULTS 5

sFigure 1. Estimated correlations among harmonized variables in the HSE and the UK Census Microdata 6

sFigure 2. Weighted and unweighted genome-wide analyses: number of genome-wide variants 7

sFigure 3. Weighted and unweighted genome-wide analyses: SNP effects 8

sFigure 4. Autosomal genome-wide association analyses on biological sex 9

sFigure 5. Genome-wide association study on UKBB participation - QQ plot 10

sFigure 6. SNP heritability estimates in weighted (wGWA) and standard genome-wide (GWA) analyses 11

sFigure 7. Genetic correlation estimates from weighted and standard genome-wide analyses 12

sFigure 8. Effect of participation bias on exposure-outcome associations obtained from Mendelian Randomization 13

SREFERENCES 14

### sMethods

#### Coding of variables included from the UKBB and HSE

| Variable | UKBB | HSE | Coding | H | G |
|---|---|---|---|---|---|
| Frequency of alcohol use | About how often do you drink alcohol? (ID: 1558) | How often have you had an alcoholic drink of any kind during the last 12 months? | 0=never, 1=few times/year, 2=few times/year, 3=monthly, 4=once or twice/week, 5=three or four days/week, daily | X | |
| Weekly alcohol use | In an average week, how many beer/cider/champagne/ wine/spirits/other alcohol) would you drink? (ID: 1588, 1578, 1608, 5364, 1568, 1598) | | continuous | | X |
| Physical activity | Number of days/week of vigorous physical activity 10+ minutes (ID:904) | | continuous | | X |
| Sex | Sex of participant | Sex of participant | 0=Male/1=Female | X | X |
| Age | Age of participant | Age of participant | Continuous | X | |
| Years of education | At what age did you complete your continuous full-time education? (ID: 845) [note: Individuals with a University degree (ID: 6138) were allocated '19 or over'] | At what age did you finish your continuous full-time education at school or college? | 14 (or under) /15/16/17/18/19 or over | X | X |
| Smoking status | Summary if the current/past smoking status of the participant (ID: 20116) | Have you ever smoked a cigarette, a cigar or a pipe? / Do you smoke nowadays? | 0=Never/1=previous/2=current | X | X |
| Vegetable intake | About how many heaped tablespoons of cooked vegetables would you eat per day? (ID: 1289) | | Continuous | | X |
| Fruit intake | About how many pieces of fresh fruit would you eat per day? (ID: 1309) | | Continuous | | X |
| Income | What is the average total income before tax received by your household? (ID: 738) | What is *your household's* income before any deductions for income tax, National Insurance, etc? | 1=18k, 2=18k-31k, 3=31k-52k, 4=52k-100k,5=>100k | X | |
| Household size | Including yourself, how many people are living together in your household? (ID: 705) | Interviewer collects the names of the people in the household | Categorical, with each category indicating the number of individuals living in the household. 7 = 7 individuals or more. | X | |
| Employment status | Which of the following describes your current situation? […] (ID: 6142) | Which of these descriptions applies to what you were doing last week? […] | 1=unemployed, 2=employed, 3=economically inactive, 4=retired | X | |
| Height | Measured using a Seca 202 device (ID: 50) | Measured during the visit | Continuous | X | X |
| Weight | Measured during the initial Assessment Centre visit (ID: 210002) | Measured during the visit | Continuous | X | |
| BMI | Derived from height and weight measures | Derived from height and weight measures | Continuous | X | X |
| BMI (categorical) | Derived from height and weight measures | Derived from height and weight measures | 1=underweight (BMI<18.5), 2=normal weight (18.5 $\leq$ BMI <25) 3=overweight (25 $\leq$ BMI <30) 4=obese (BMI $\geq$ 30), | X | |
| Non-cancer diagnoses (number) | Number of self-reported non-cancer illnesses (ID: 135) | | Continuous | | X |
| Risk taking | Would you describe yourself as someone who takes risks? (ID: 2040) | | 0=No/1=Yes | | X |
| Loneliness | Do you often feel lonely? (ID: 2020) | | 0=No/1=Yes | | X |
| Diabetes | Has a doctor ever told you that you have diabetes? (ID: 2443) | | 0=No/1=Yes | | X |
| Depression/Anxiety | Seen a psychiatrist for nerves, anxiety, tension or depression (ID: 2100) | | 0=No/1=Yes | | X |
| LDL | LDL cholesterol (ID: 30780) | | Continuous | | X |
| SBP | Systolic blood pressure, automated reading (ID: 4080) | | Continuous | | X |
| Reaction time | Reaction time (mean time to correctly identify matches) (ID: 20023) | | Continuous | | X |
| Insomnia | Do you have trouble falling asleep at night or do you wake up in the middle of the night? (ID: 1200) | | 1=Never/rarely, 2=sometimes, 3=usually | | X |

| | | | | | |
|---|---|---|---|---|---|
| Cancer | Has a doctor ever told you that you have had cancer? (ID: 2453) | | 0=No/1=Yes | | X |
| Coffee intake | How many cups of coffee do you drink each day? (ID: 1498) | | Continuous | | X |
| Urbanisation | Classification derived by combining each participant's home postcode with data generated from the 2001 census from the Office of National Statistics (ID: 20118) | Degree of urbanisation | 1=village/hamlet, 2=town/fringe, 3=urban | X | |
| Overall health | In general, how would you rate your overall health? (2178) | How is your health in general? | 1=poor; 2=fair, 3=good | X | |

H=used for harmonization and included in the model predicting participation probability; G=included as outcome in genome-wide analyses

#### Coding of variables harmonized across the UKBB, HSE and the UK Census Microdata

| Variable | UKBB | HSE | Census | Coding |
|---|---|---|---|---|
| Sex | Sex of participant | Sex of participant | Sex of participant | 0=Male/1=Female |
| Age | Age of participant | Age of participant | Age of participant | 40-44 / 45-49 / 50-54 / 55-59 / 60-64 / 65-69 |
| Years of education | At what age did you complete your continuous full-time education? (ID: 845) [note: Individuals with a University degree (ID: 6138) were allocated '19 or over'] | At what age did you finish your continuous full-time education at school or college? | Level of highest qualifications:<br>14 (or under)=No academic or professional qualifications<br>15=Other (vocational/foreign/outside UK quals)<br>15=Level 1 (0-4 GCSE, O level, or equivalents)<br>16=Level 2 (5+ GCSE, O level, 1 A level, or equivalents)<br>17=Apprenticeship<br>17=Level 3 (2+ A levels, or equivalents)<br>19 (or over)=Level 4+ (degree, postgrad, professional quals) | 14 (or under) /15/16/17/18/19 or over |
| Employment status | Which of the following describes your current situation? […] (ID: 6142) | Which of these descriptions applies to what you were doing last week? […] | Employment Status based on the International Labour Office (ILO) definition. | 1=unemployed, 2=employed, 3=economically inactive, 4=retired |
| Overall health | In general, how would you rate your overall health? (2178) | How is your health in general? | Self-reported health | 1=poor; 2=fair, 3=good |

#### Genotyping, imputation and quality control in the UK Biobank

488,377 UKBB participants were genotyped, using either the Applied Biosystems UK BiLEVE Axiom Array (807,411 markers for 49,950 participants) or the Applied Biosystems UK Biobank Axiom Array (825,927 markers for 438,427 participants). Poor quality samples were identified using the metrics of missing rate and heterozygosity computed using a set of 605,876 high quality autosomal markers that were typed on both arrays. Imputation was performed using IMPUTE4 with the Haplotype Reference Consortium (HRC) UK10K and the 1000 Genomes Phase 3 dataset as the main imputation reference panels. Detailed genotyping, imputation and quality control (QC) procedures have previously been described[1]. Additional quality control filters for genome-wide analyses were applied to select participants (i.e., exclusion of related individuals, exclusion of non-White British ancestry based on principal components, high missing rate and high heterozygosity on autosomes) and genetic variants (Hardy–Weinberg disequilibrium $P>1\times10^{-6}$, minor allele frequency>1% and call rate>90%).

### sResults

*Probability weighted genome-wide association analyses on UK Biobank traits*

Among all genome-wide hits (1690, with $p<5\times10^{-8}$), overestimation was more common (420 SNPs, 24.85% of all genome-wide SNPs) than underestimation (290 SNPs, 17.16% of SNPs). Change in direction of SNP effects was rare, as was the case for only one of all 1690 identified SNPs (rs2163971 on smoking status). However, the effects were only significant in standard GWA ($\hat{\beta}$=0.011, p=1.13e-09) but not weighted GWA ($\hat{\beta}_w$=-0.001, $p_w$=0.776).

*Weighted SNP heritability and genetic correlation estimates*

A number of the assessed trait-pairs were significantly underestimated or overestimated as a result of participation bias. Change in direction of genetic correlations as a result of participation bias was less present. While a number of genetic correlations showed opposite signs between *rg* and $rg_w$ (17 out of the 153 assessed trait pairs), most of these $rg_{DIFF}$ ($rg$- $rg_w$) were not significantly different ($p<0.05$). For example, the largest $rg_{DIFF}$ with opposite signs in *rg* and $rg_w$ was present for *rg*(depression/anxiety, vegetable intake) [*rg*=0.19; *p*=4.3e-05 versus $rg_w$=-0.12; *p*=0.45, FDR-corrected *p*-value for $rg_{DIFF}$ = 1] and *rg*(number of illnesses, vegetable intake) [*rg*=0.19; *p*=7e-07 versus $rg_w$=-0.01; *p*=0.9, FDR-corrected *p*-value for $rg_{DIFF}$ = 1].

*Effect of participation bias on Mendelian Randomization estimates*

Of all exposure-outcome associations tested (k=234), 14 (6%) estimates were either overestimated or underestimated. Significant ($p_{FDR}<0.05$) differential effects were only present for two of the exposure-outcome associations tested (education on BMI; smoking status on fruit consumption). There was little evidence of bias resulting in changes in direction of MR estimates. The largest difference between $\hat{\alpha}$ and $\hat{\alpha}_w$ resulting from opposite effects was present for fruit intake on LDL cholersterol ($\hat{\alpha}$=0.03; *p*=0.83 versus $\hat{\alpha}_w$=-0.12; *p*=0.47) and smoking status on physical activity ($\hat{\alpha}$=0.07; *p*=0.091 versus $\hat{\alpha}_w$=-0.04; *p*=0.45).

sFigure 1. Estimated correlations among harmonized variables in the HSE and the UK Census Microdata

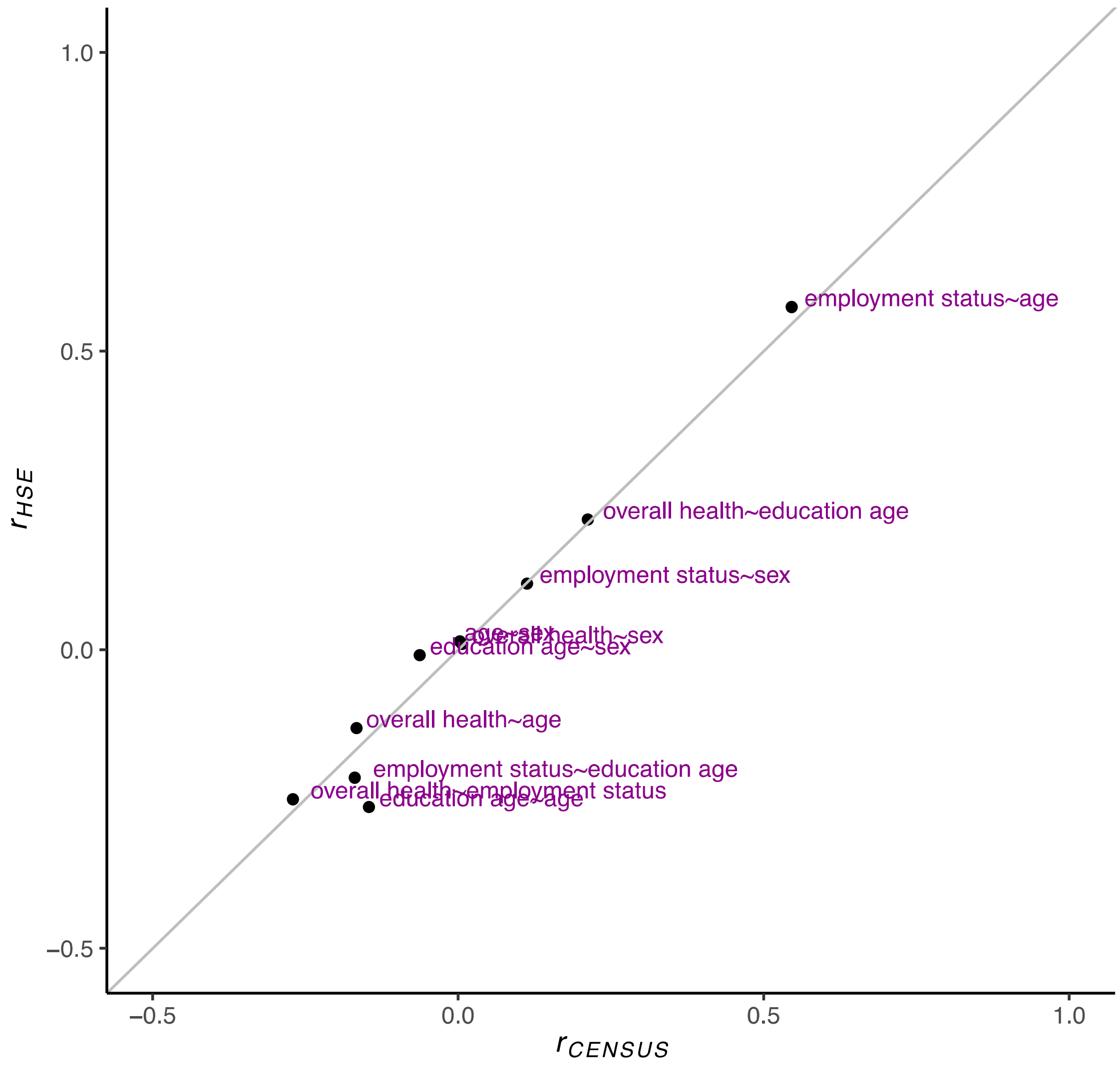

sFigure 2. Weighted and unweighted genome-wide analyses: number of genome-wide variants

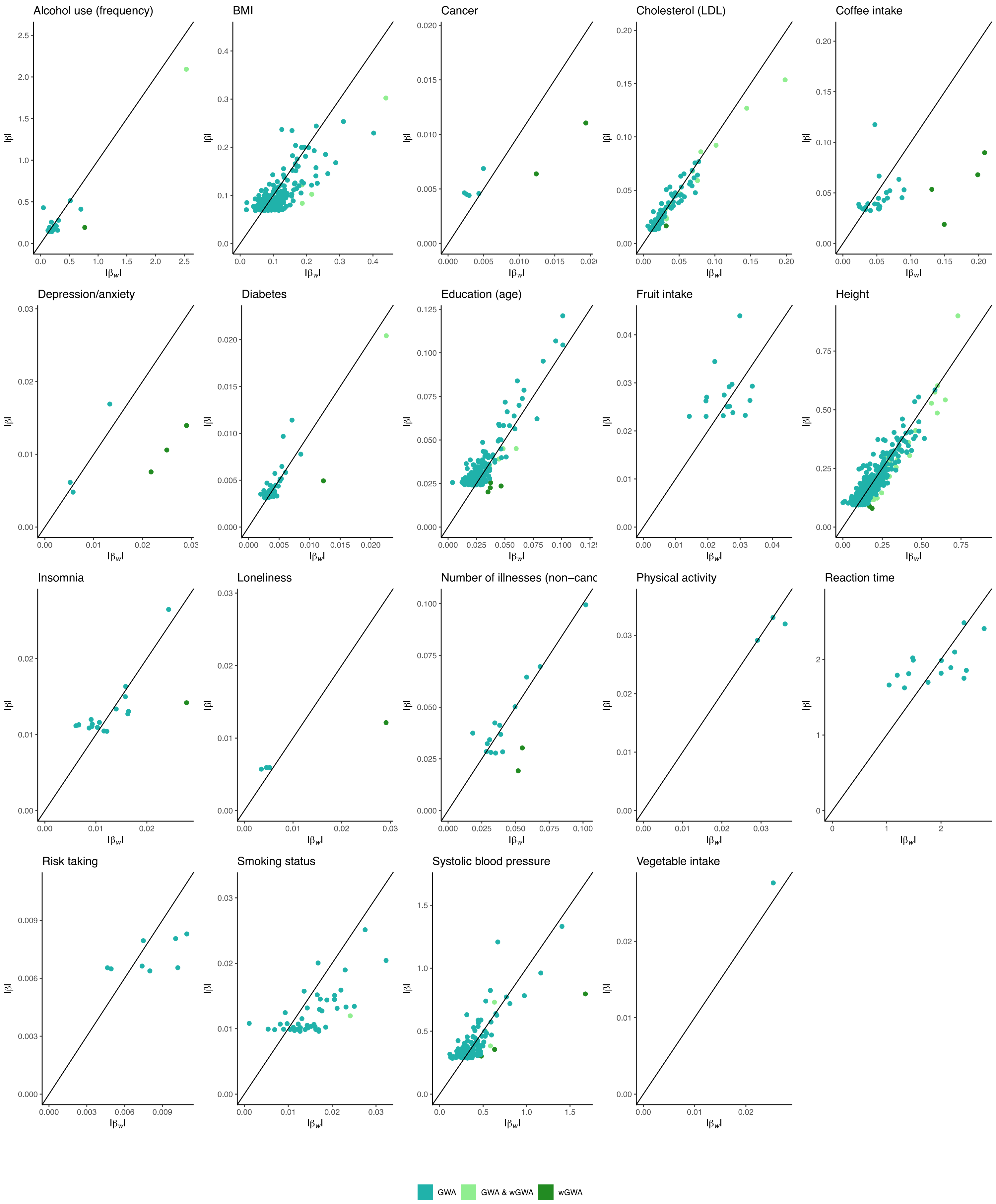

### sFigure 3. Weighted and unweighted genome-wide analyses: SNP effects

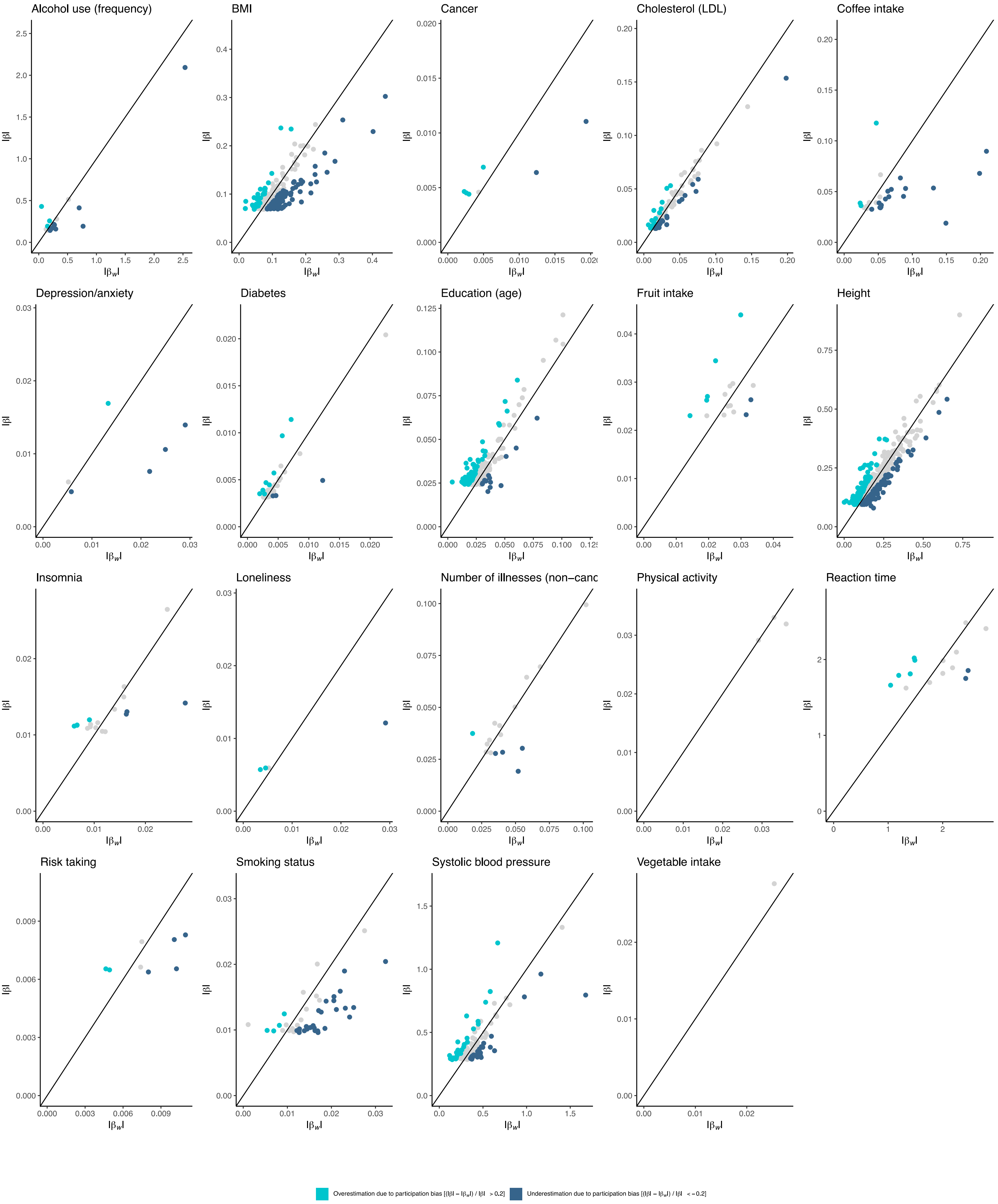

sFigure 4. Autosomal genome-wide association analyses on biological sex

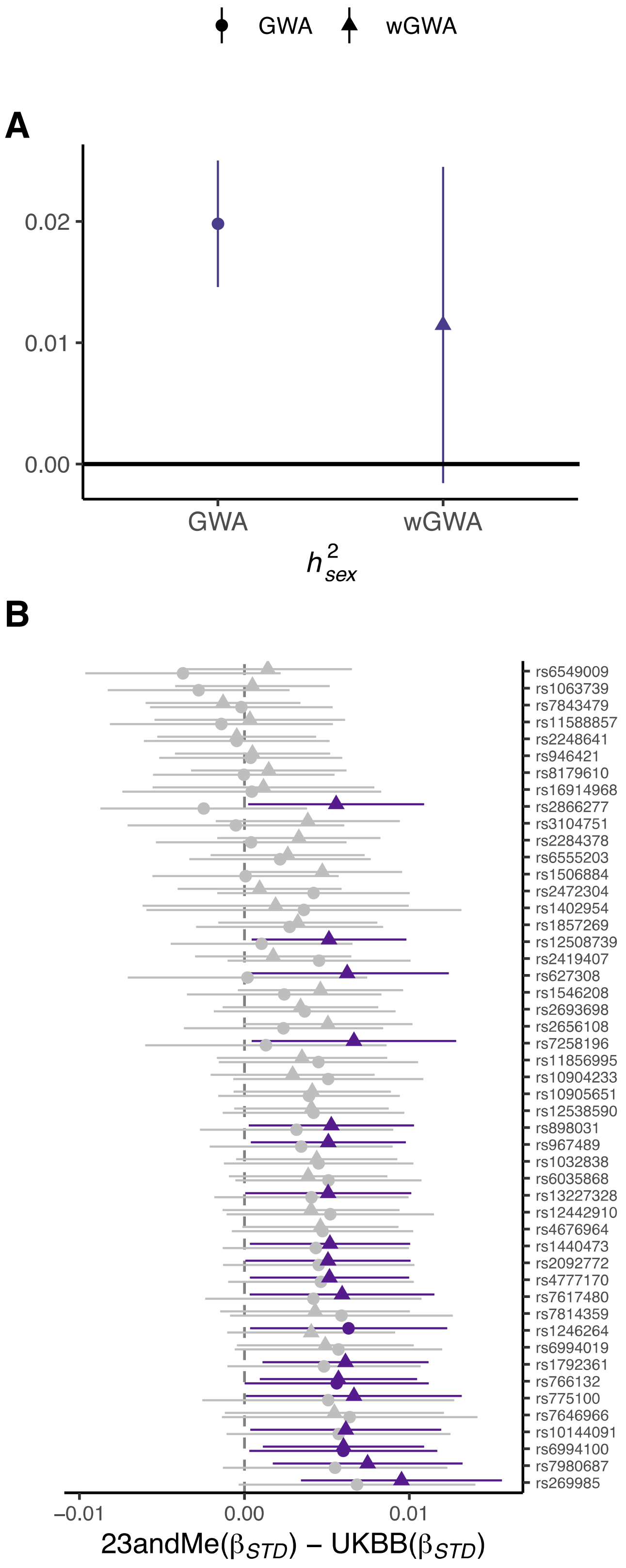


Panel (**A**) displays the SNP heritability estimates of sex-associated variants obtained from standard GWA and wGWA. Panel (**B**) displays the effects of 49 autosomal variants on sex, comparing estimates obtained from standard GWA and wGWA to estimates obtained from an independent sample of >2,400,000 volunteers.

sFigure 5. Genome-wide association study on UKBB participation - QQ plot

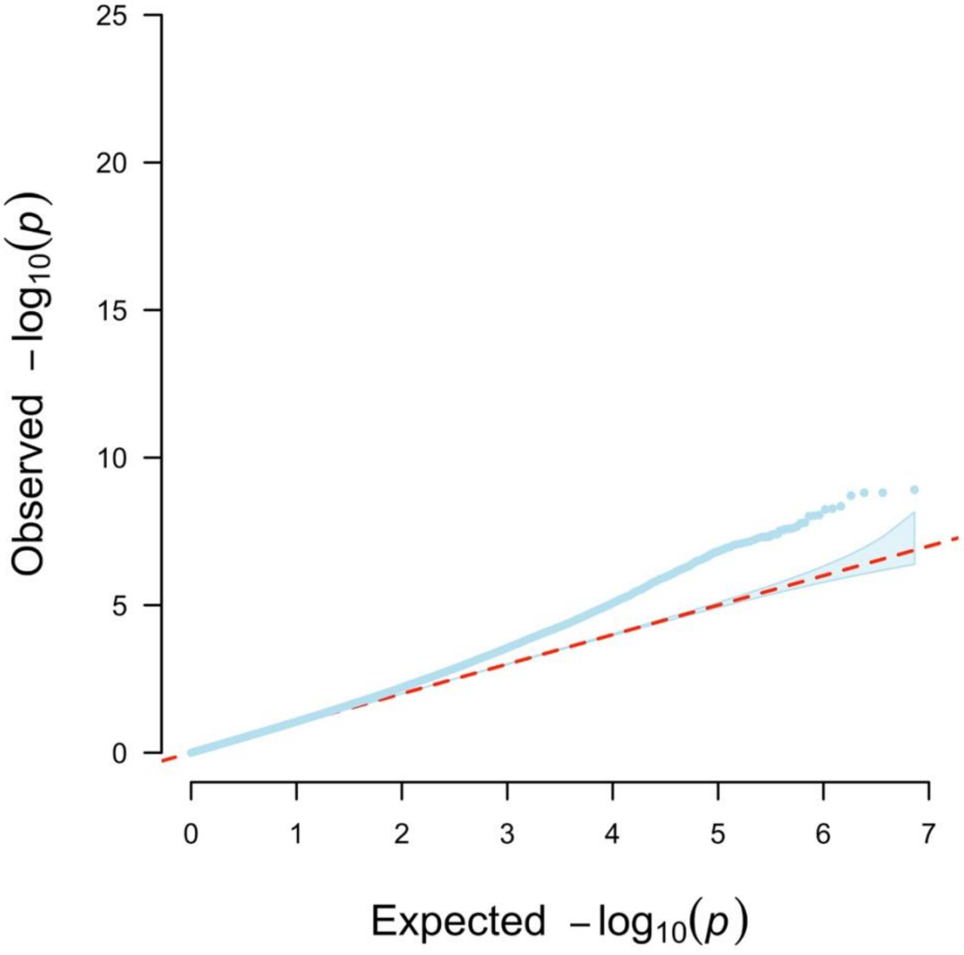

sFigure 6. SNP heritability estimates in weighted (wGWA) and standard genome-wide (GWA) analyses

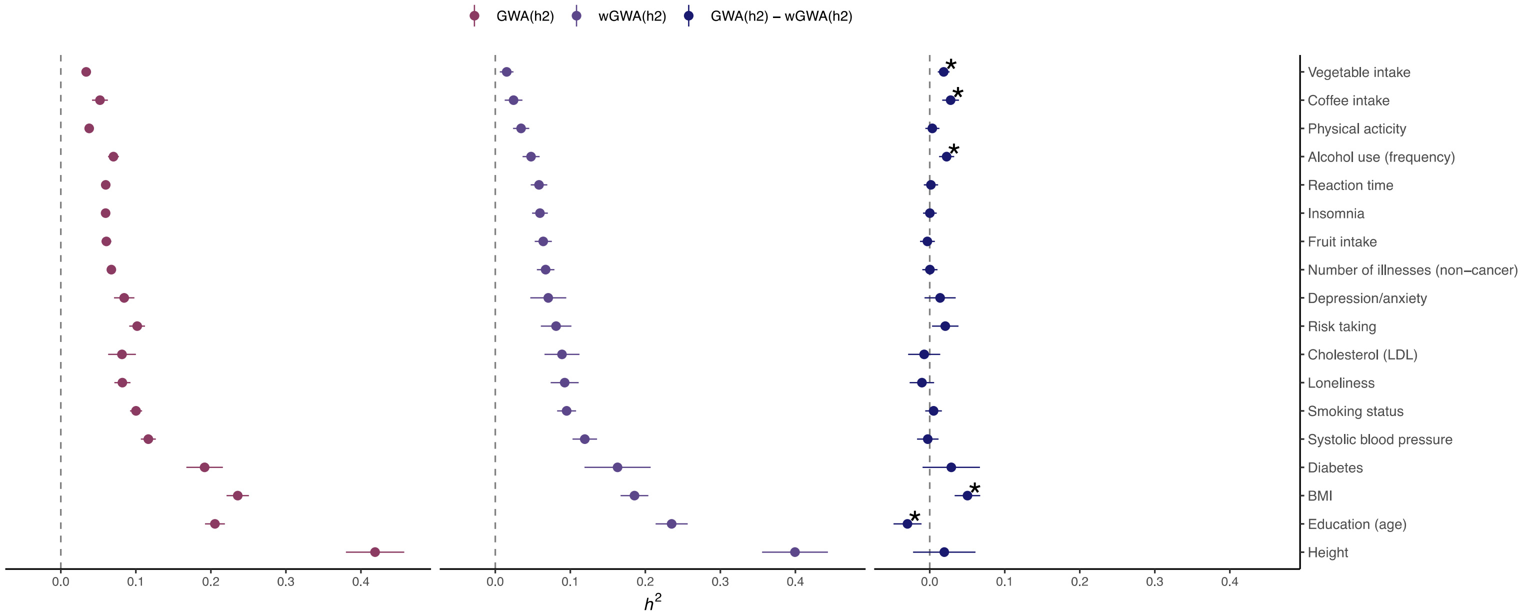


LDSC heritability ($h^2$) estimates, obtained using the output from standard (unweighted) GWA analyses [GWA(h2)] and probability weighted GWA [wGWA(h2)]. The right panel displays the differences in SNP heritability between standard and weighted GWA ($h^2$ - $h_w^2$). (*) Estimates showing significant differences ($p_{FDR}$<0.05)

#### sFigure 7. Genetic correlation estimates from weighted and standard genome-wide analyses

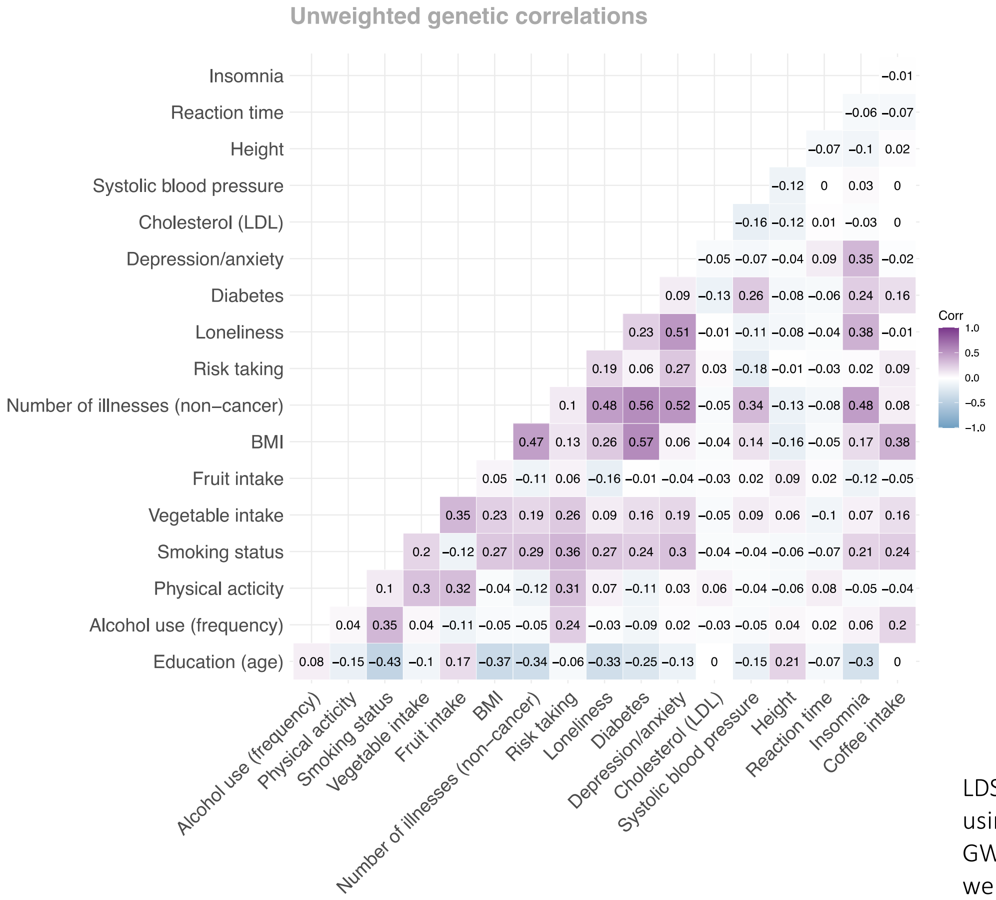


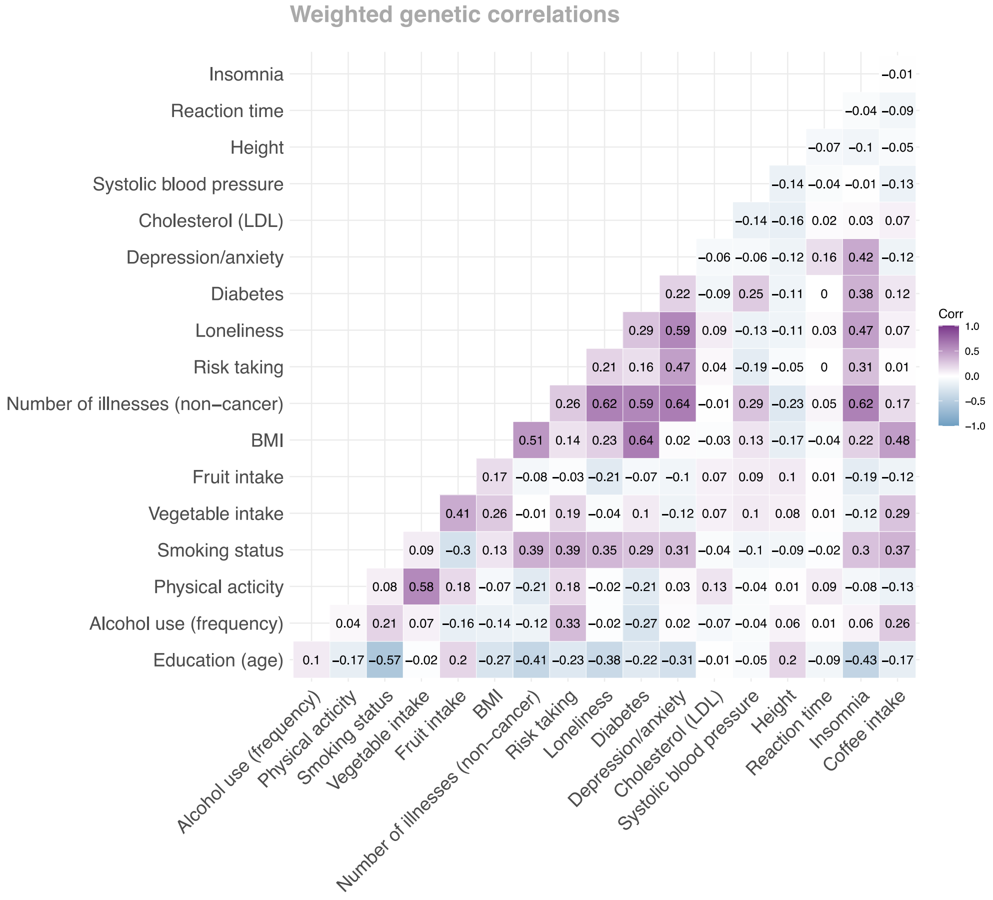


LDSC genetic correlations estimates, obtained using the output from standard (unweighted) GWA analyses (upper panel) and probability weighted GWA (lower panel)

### sFigure 8. Effect of participation bias on exposure-outcome associations obtained from Mendelian Randomization

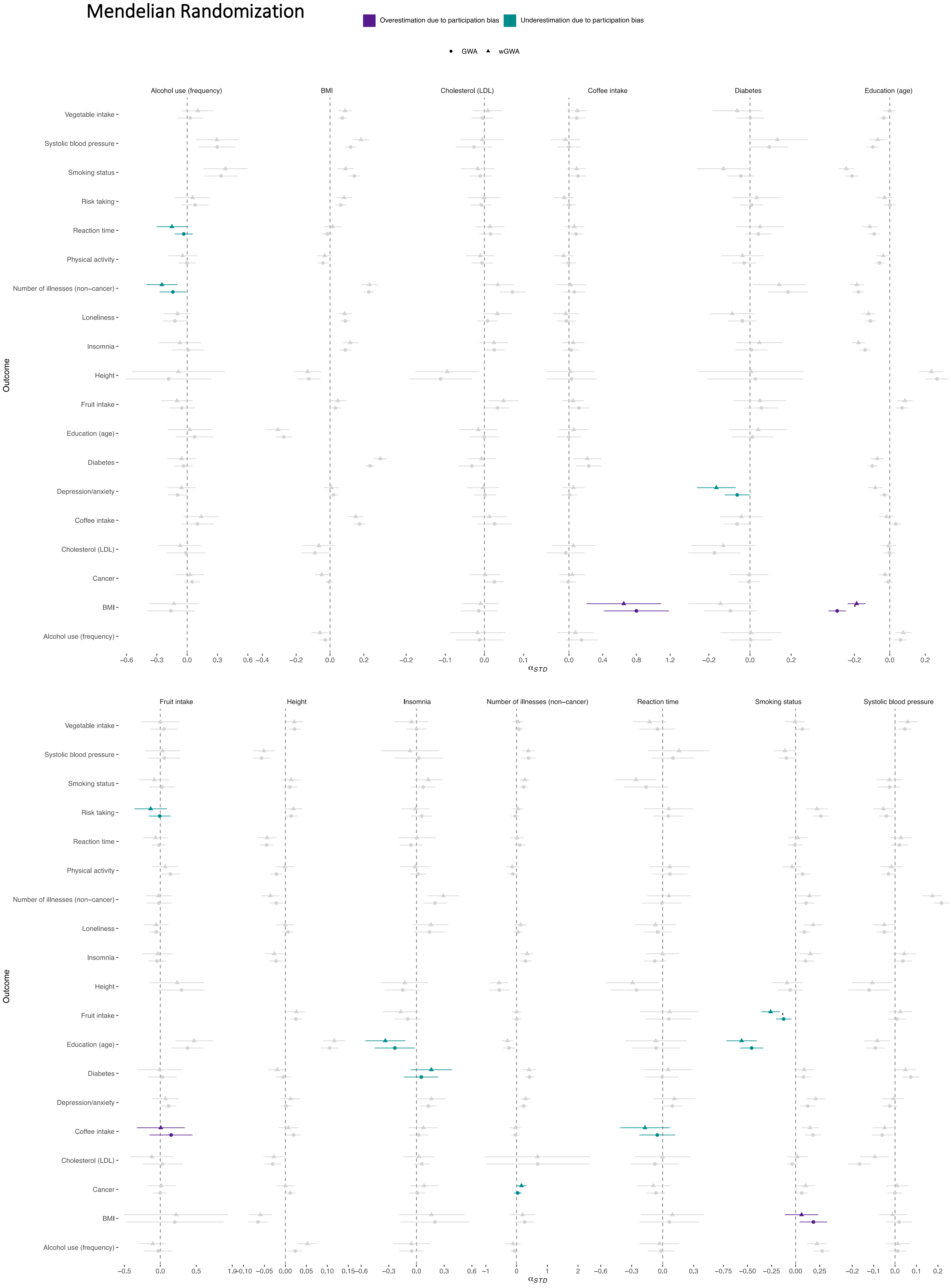

### sReferences

1. Bycroft C, Freeman C, Petkova D, et al. The UK Biobank resource with deep phenotyping and genomic data. *Nature*. 2018;562(7726):203-209. doi:10.1038/s41586-018-0579-z
